# Rapid, one-step generation of biallelic conditional gene knockouts

**DOI:** 10.1101/056549

**Authors:** Amanda Andersson-Rolf, Roxana C. Mustata, Alessandra Merenda, Sajith Perera, Tiago Grego, Jihoon Kim, Katie Andrews, Juergen Fink, William C. Skarnes, Bon-Kyoung Koo

## Abstract

Loss-of-function studies are key to investigate gene function and CRISPR technology has made genome editing widely accessible in model organisms and cells. However, conditional gene inactivation in diploid cells is still difficult to achieve. Here, we present CRISPR-FLIP, a strategy that provides an efficient, rapid, and scalable method for bi-allelic conditional gene knockouts in diploid cells by co-delivery of CRISPR/Cas9 and a universal conditional intron cassette.

## Main text

Analysing gene function is a crucial step in our understanding of normal physiology and disease pathogenesis. In cell-based models, loss-of-function studies require inactivation of both copies of the gene. Prior to the development of site-specific nucleases, gene knockouts in cell lines were achieved by loss-of-heterozygosity^1^ or serial gene targeting approaches^2^. The development of site-specific nucleases, such as zinc finger nucleases, has greatly facilitated functional studies in cells due to the fact that both copies of a gene can be efficiently inactivated in a single step^3^. Recently, the CRISPR-Cas9 gene editing technology^4–7^ has become the tool of choice for gene knockout studies due to its simplicity and robustness. Cas9 nuclease is an RNA-guided nuclease that is highly efficient in inducing a double-strand break (DSB) at a genomic site of interest. These DSBs can be repaired by error-prone non-homologous end joining (NHEJ) to generate gene-inactivating mutations or, in the presence of a donor template, the DSBs can be repaired by homology-directed repair (HDR) to generate more precise and complex alleles^8^. While simple constitutive knockouts are useful and informative, in many cases it is desirable to engineer conditional loss-of-function models, particularly for genes essential for cell viability or embryonic development. Here, we describe a simplified, one-step method for engineering conditional loss-of-function mutations in diploid cells. We developed a novel, invertible drug selection cassette, FLIP, for high-efficiency nuclease-assisted targeting in cells. We are able to recover biallelic events from a single round of gene targeting and screening. As proof-of-principle, we applied our method to conditionally inactivate both essential and nonessential genes in mouse and human cells.

Existing methods for engineering conditional mutations in cultured cells^9–12^ rely on the inclusion of a drug selection cassette that must be removed in a second step to ensure proper expression of targeted conditional allele (see Supplementary Fig 1). These methods were not designed for the generation of conditional loss-of-function models in a single step, particularly where the target gene is essential for cell growth or viability. To overcome these limitations, our strategy combines an invertible intronic cassette (FLIP), similar to COIN^12^, with high efficiency Cas9-assisted gene editing. Critically, the non-mutagenic orientation of the FLIP cassette expresses the puromycin resistance gene (puroR) to select for correct nuclease-assisted targeting into the exon of one allele and simultaneous enrichment of cells that inactivate the second allele by nuclease-mediated NHEJ (Fig 1a). Upon exposure to Cre recombinase the FLIP cassette is inverted to a mutagenic configuration that activates a cryptic splice acceptor and polyadenylation signal (pA) and disrupting the initial splicing acceptor resulting in the complete loss of gene function (Fig. 1b and Supplementary Fig. 2a). In contrast to COIN which requires the removal of the drug selection cassette, our FLIP cassette permits the generation of conditional mutant cells in one step.

**Figure 1.**
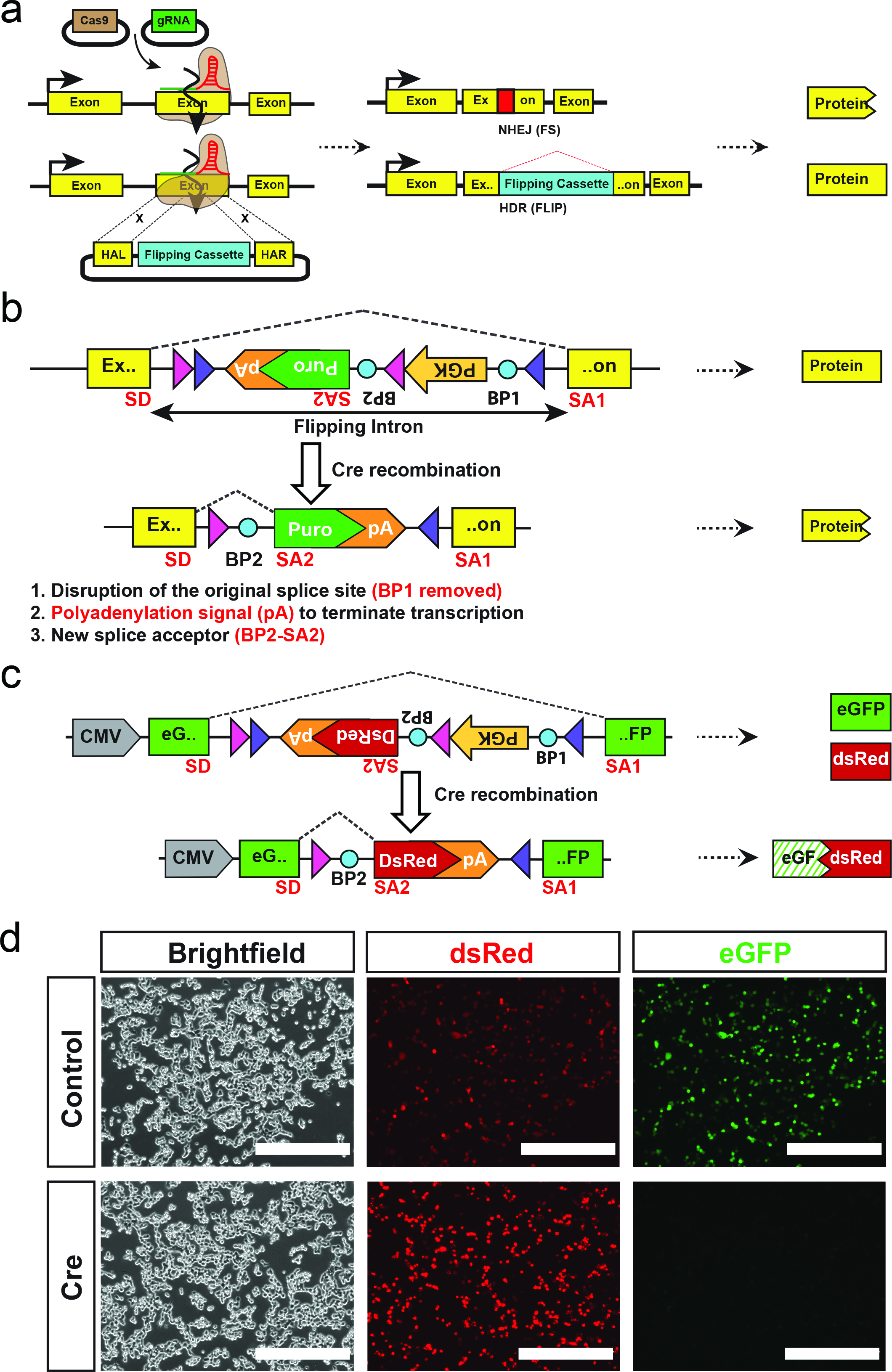
FLIP cassette strategy for bi-allelic conditional gene modification
(a) Schematic drawing of the FLIP cassette strategy for bi-allelic conditional gene modification. The Cas9 nuclease is directed to the genomic site of interest by the gRNA where it generates a double stranded break (DSB, left panel). This DSB can be repaired by non-homologous end joining (NHEJ) generating insertions/deletions (indels) or homology directed repair (HDR, right panel). In the latter, the donor plasmid is used as template for precise correction of the DSB and thus facilitating insertion of FLIP cassette in the genome. Bi-allelic conditional gene modification is achieved when one allele is repaired via NHEJ generating a frame shift mutation and the other through HDR, resulting in FLIP cassette insertion.
(b) The design of the FLIP cassette. The FLIP cassette contains several elements: i) a reporter or resistance gene which expression is controlled by a promoter and polyadenylation signal (initially in the antisense direction) ii) two pairs of loxP sites for Cre mediated recombination iii) splicing donor and acceptor sites (including two branching points) for spliceosome recognition and intron excision. The FLIP cassette is flanked with left and right homologous arms, each arm less than 1kb, generating a targeting vector. Following Cre recombination the cassette is inverted (flipped) and a new splicing configuration is activated. This results in inactivation of the gene via three rearrangements: the old splice site is disrupted (BP1 removed), the inversion of the pA and new splice (BP2) signal into the sense direction leads to termination of transcription. SD - splice donor, SA1, SA2 - splice acceptor, loxP sites - pink and purple triangles -, BP1, BP2 (blue circles) - branching point, pA - polyadenylation signal.
(c) The FLIP cassette containing a DsRed reporter gene is inserted into the cDNA of eGFP as an artificial intron and transfected in HEK 293T cells.
(d) Following insertion, the cassette functions as an intron and does not disrupt the expression of the eGFP cDNA. Hence, both eGFP and DsRed proteins are expressed (top row). After Cre treatment the eGFP expression is disrupted, and only DsRed expression is maintained (bottom row). Scale bar 400μm.

Initially, to test the functionality of our intronic FLIP cassette, we inserted a FLIP cassette variant containing a dsRed2 reporter in place of puroR into a CMV-eGFP (enhanced green fluorescent protein) expression plasmid (Fig. 1c). Following transient transfection of HEK293 cells, both green and red fluorescence was observed, demonstrating that insertion of the FLIP cassette in the non-mutagenic orientation is inert (Fig. 1d). Cre recombinase-mediated FLIP cassette inversion resulted in loss of eGFP expression, showing conditional inactivation of eGFP expression in the inverted, mutagenic orientation (Fig. 1c,d).

Next, we employed CRISPR/Cas9 endonuclease in mouse embryonic stem cells (mESCs) to introduce the puroR FLIP cassette into one allele of β-catenin (*Ctnnb1*) via HDR and to simultaneously induce a frameshift mutation by NHEJ in the second β-catenin allele (Fig. 1a, Fig. 2a). β-catenin is an important gene for the morphology and efficient self-renewal of mESCs^13,14^. A donor vector containing the puroR FLIP cassette inserted in exon 5 of β-catenin and flanked by ˜1 kb homology arms was transfected into mESCs with Cas9 and gRNA expression plasmids. Following selection in puromycin, drug-resistant colonies were genotyped by PCR to confirm correct integration of the FLIP cassette and then assayed for NHEJ events in the second allele by Sanger sequencing (Fig. 2b,c, Supplementary Fig. 2b). From 64 clones, 14 clones (21.9%) were correctly targeted, among which 4 clones carried a frame-shift mutation in the second allele (Table 1). The recovery of β-catenin compound mutant clones (FLIP targeted/NHEJ frameshift; FLIP/-) with wildtype morphology strongly suggests that the insertion of the FLIP cassette does not disrupt the function of β-catenin in the non-mutagenic orientation. Upon expression of Cre recombinase in FLIP/- clones, we observed a loss of β-catenin expression in cells (Fig. 2d, e). Moreover, compared to control (FLIP/+) cells treated with Cre recombinase, the FLIP/- cells became scattered and lost their dome-like morphology (Fig 2f).

**Figure 2.**
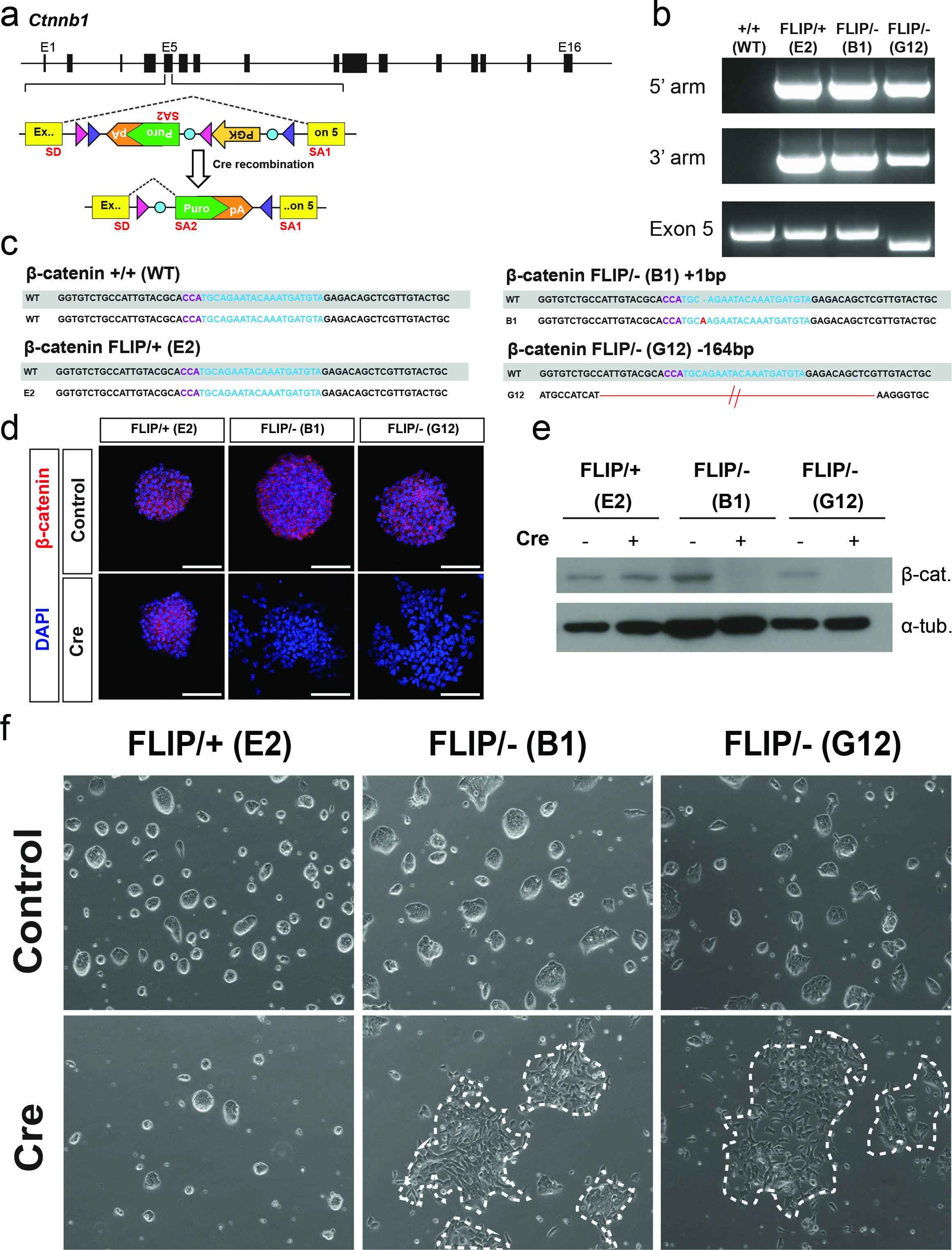
Insertion of the FLIP cassette in the endogenous *Ctnnb1* gene of mouse embryonic stem cells.
(a) The FLIP cassette containing a resistance gene is inserted into the 5th exon of *Ctnnb1.* SD - splice donor, SA - splice acceptor, pink and purple triangles - loxP site, BP - branching point.
(b) PCR detection of FLIP cassette insertion in the *Ctnnb1* locus. Correctly targeted clones E2, B1, and G12 are positive for 5′ and 3′arm genotyping PCR reactions (for genotyping strategy see Fig. S2b). Exon 5 PCR detects the remaining allele. The clones (E2, B1 and G12) are correctly targeted.
(c) Insertions/deletions in the non-targeted allele were identified by Sanger sequencing. Clone B1 has a 1 base pair (bp) insertion and clone G12 has a 164bp deletion. gRNA and PAM recognition sequences are represented in blue and purple respectively. The predicted wild type (WT) sequence is shown in a grey box and the actual sequence of the second allele (not having a FLIP cassette inserted) is aligned underneath. (d, e) Detection of β-catenin protein by immunofluorescence (d) and western blotting (e) before and after Cre transfection. This confirms the loss of β-catenin on protein level for the FLIP/clones, B1 and G12 following Cre treatment.
(f) Representative bright field images of the ESC clones before (top) and after (bottom) Cre transfection. B1 and G12 clones displays altered phenotype due to *Ctnnb1* gene inactivation.

**Table 1.** Summary of targeting using CRISPR-FLIP strategy.

| Targeted gene | Total No. of clones | No. of clones with correct insertion of FLIP | Targeting efficiency (%) | FLIP/+ clones | FLIP/- clones (total) | FLIP/- in-frame | FLIP/- out of frame | Misc. (mixed, not sequence confirmed clones and potentially empty wells) |
| --- | --- | --- | --- | --- | --- | --- | --- | --- |
| *Apc* | 89 | 27 | 30.3% | 13 | 6 | 0 | 6 | 8 |
| *Ctnnb1* | 64 | 14 | 21.9% | 7 | 6 | 2 | 4 | 1 |
| *Esrrb* | 192 | 52 | 27.1% | 25 | 4 | 1 | 3 | 23 |
| *Nfx1* | 96 | 19 | 19.8% | 4 | 3 | 1 | 2 | 12 |
| *Sox2* | 192 | 58 | 30.2% | 37 | 3 | 2 | 1 | 18 |
| *Tcf7l2* | 96 | 39 | 40.6% | 17 | 14 | 3 | 11 | 8 |
| *Trim13* | 96 | 21 | 21.9% | 4 | 4 | 0 | 4 | 13 |
| *Trim37* | 96 | 28 | 29.2% | 4 | 8 | 3 | 5 | 16 |
| *ARID1A* (HEK293) | 192 | 18 | 9.4% | 6 | 3 | 0 | 3 | 9 |
| *TP53* (HEK293) | 104 | 42 | 40.4% | 2 | 8 | 3 | 5 | 32 |
| *TP53* (hiPSC) | 32 | 24 | 75.0% | 12 | 4 | 2 | 2 | 8 |

To test if the CRISPR-FLIP technology is widely applicable, we additionally targeted *Apc, Esrrb, Nfx1, Sox2, Tcf7l2, Trim13,* and *Trim37* in mESCs; *TP53* and *ARID1A* in human HEK293 cells; and *TP53* in human induced pluripotent stem cells (Supplementary Fig. 3-7). The FLIP intron targeting efficiency ranged from 19.8% to 40.6% in mESCs (Table 1). For all genes, FLIP/− clones were obtained (Table 1, Supplementary Fig. 4-7). The conditional inactivation of gene expression was confirmed by Western blot and immunofluorescence for Esrrb and Sox2 (Supplementary Fig. 4). We conclude that the FLIP conditional strategy is efficient and can be applied widely for conditional loss-of-function studies in various mammalian cells that are amenable to Cas9-assisted gene targeting.

Our strategy requires the presence of a CRISPR site overlapping or nearby the insertion site of the FLIP cassette, imposing constraints on the exons than can be targeted. To maximize the potential for a null mutation, the target exon must be common to all transcripts and lie within the first 50% of the protein-coding sequence. Additionally, based on the minimum size of mammalian exons (50 bp)^15^, we set the size of the split exons to be at least 60 bp. Finally, for optimal splicing, we chose insertion points that match the consensus sequence for mammalian splice junctions (minimally MAGR (^A^/_C_AG/Pu))^16^. Using this set of rules, we used bioinformatics to estimate the number of suitable FLIP insertion sites in the protein-coding genes in the mouse and human genomes. Our bioinformatics analysis revealed 1,171,712 FLIP insertion sites and corresponding gRNA binding sites covering 16,460 genes in the mouse genome and 1,171,787 FLIP insertion sites and corresponding gRNA binding sties covering 15,177 genes in the human genome. (Supplementary Table 1).

Here we present the FLIP technology, a method that allows one-step generation of bi-allelic conditional gene modifications using only a single gRNA, Cas9 and a simple donor vector. Compared to the conventional strategies for the generation of conditional alleles, the FLIP cassette, when combined with the CRISPR/Cas9, enables highly efficient bi-allelic conditional gene modification in a single round of gene targeting without the need to remove the drug selection cassette. The FLIP targeting vectors only require short homologous arms (less than 1 kb) which makes the assembly of targeting vectors easy and scalable. The FLIP cassette is invariable and can be generically applied to any gene, including non-coding RNA genes.

## Methods

### dsRed FLIP cassette inserted in the eGFP cDNA

The FLIP cassette inserted in the middle of eGFP and containing a dsRed2 reporter gene was synthesized and ordered from GenScript. The split eGFP cDNA and the FLIP cassette were cloned into the mammalian expression vector pCDNA4TO (Invitrogen) using BamHI (R0136S, NEB) and XhoI (R0146S, NEB) (for pre-recombined form). The vector was subsequently transformed into Cre expressing bacteria (A111, Gene bridges) to generate the Cre-recombined form. Correct clones were confirmed with restriction digest BamHI (R0136S, NEB) and XhoI (R0146S, NEB) and Sanger sequencing.

### FLIP cassette containing selection marker genes

The FLIP cassette was PCR amplified and cloned into Pjet1.2 vector (ThermoFisher Scientific, K131). Replacement of dsRed was done through restriction digest excision using EcoRI (R3101S, NEB) and Acc65l (RO599S, NEB) followed by insertion of PCR amplified selection marker genes using Infusion cloning (638909, Clontech). The FLIP cassette including selection marker gene was then transferred to the vector pUC118 (3318, Clontech) using the restriction enzymes SacI (R0156S, NEB) and PstI (R0140S, NEB) and Mighty cloning (6027, Takara).

### Addition of homologous arms to the FLIP cassette - FLIP targeting vector generation

Homologous arms around an intron insertion site were amplified by high fidelity Phusion DNA polymerase (M0530S, NEB). After PCR product purification, both homologous arms and FLIP cassette-containing vector were mixed with the type II restriction enzyme SapI and T4 DNA ligase (M0202T, NEB). After 25 cycles of 37°C and 16°C, the reaction mixture was directly used for E.Coli transformation. DNA was extracted (27106, Qiagen) and analysed with restriction digest to identify correctly assembled FLIP donor vectors.

### Cas9 and gRNA plasmids

Human codon optimized Cas9 (41815, Addgene) and empty gRNA vector (41824, Addgene) were obtained from Addgene.

### Cell culture

***HEK293 cells.*** Human embryonic kidney 293 cells were cultured in media consisting of DMEM, high glucose (11965092, Thermofisher Scientific) supplemented with 10% foetal bovine serum (Thermofisher Scientific), 1x penicillin-streptomycin according to the manufacturer’s recommendation (P0781, Sigma).

***Embryonic stem cells (ESCs).*** Murine E14 Tg2a embryonic stem (mES) cells were cultured feeder-free on 0.1% gelatin-coated dishes in serum+LIF+2i (Chiron and PD03) composed of GMEM (G5154, Sigma), 10% foetal bovine serum (Gibco), 1x non-essential amino acids according to the manufacturer’s recommendation (11140, Thermofisher Scientific), 1mM sodium pyruvate (113-24-6, Sigma), 2mM L-glutamine (25030081, Thermofisher Scientific), 1x penicillin-streptomycin according to the manufacturer’s recommendation (P0781, Sigma) and 0.1mM 2-mercaptoethanol (M7522, Sigma), 20ng/ml murine LIF (Hyvonen lab, Cambridge), 3 μM CHIR99021 and 1 μM PD0325901 (Stewart lab, Dresden). BOBSC^17^ human induced pluripotent stem (hiPS) cells were cultured feeder-free on dishes coated with Synthemax II (3535, Corning) in TeSR-E8 media (05940, Stem Cell Technologies). ESCs were kept in a tissue culture incubator at 37°C and 5% CO_2_. Cells were split in a 1:10 – 1:15 ratio every 3-4 days depending on confluence.

### Cell Transfections

For targeting of mouse ESCs 1×10^6^ cells were collected and resuspended in magnesium and calcium free phosphate buffered saline (D8537, Sigma). A total of 50μg of DNA consisting of the targeting vector, Cas9 and gRNA in a 1:1:1 ratio were added to the cells and then transferred to a 4mm electroporation cuvette (Biorad). Electroporation was performed using the Biorad Gene Pulser XCell’s (165–2660, Biorad) exponential program and the following settings: 240V, 500uF, unlimited resistance. For targeting of human iPS cells, 2×10^6^ cells were dissociated with Accutase (SCR005, Millipore) and resuspended in nucleofection buffer (Solution 2, LONZA). A total of 12 μg of DNA consisting of 4 μg Cas9 plasmid, 4 μg of each gRNA plasmid and 4 μg of targeting vector was added to the cells and transferred to a 100 μl nucleofection cuvette (LONZA). Nuclefection was performed with the AMAXA Human Nucleofector Kit 2 (LONZA Cat # VPH-5022) using the B-016 program. The cells were plated and cultured for 1 day in TeSR-E8 media containing ROCK inhibitor (Y-27632, Stem Cell Technologies) to promote survival of transfected cells.

For targeting of HEK293 cells, the cells were cultured until they reached 50–60% confluence. A total of 8μg of DNA consisting of targeting vector, Cas9 and gRNA in a 1:1:1 ratio was transfected using Lipofectamine 2000 (11668019, Invitrogen) according to the manufacturer's instructions.

### Cre Transfection

1μg of pCAG-Cre-Puro plasmid vector and 3μl of Lipofectamine2000 (11668019, Invitrogen) were mixed according to the manufacturer's protocol, applied to 200.000 cells/ 6-well and incubated overnight. Media was refreshed the following morning.

### Western Blot

Following transfection ESCs were cultured for 2–5 days and then lysed in buffer containing complete protease-inhibitor cocktail tablets (11697498001, Roche) and centrifuged at 13000rpm for 15min at 4°C. Protein concentration was measured with Bradford assay (5000204, Biorad) and equal amounts were loaded on a 10% acrylamide gel and run at 120V for 1.5–2hrs. The proteins were subsequently transferred to an Immobilon-FLPVDF 0.45μm membrane (IPFL00010, Millipore) at 90V for 1hr 15min. The following primary antibodies and dilutions were used to detect the indicated proteins: Rabbit monoclonal antibody against β-catenin (1:1000, 8480S, Cell Signaling), mouse monoclonal against alpha Tubulin antibody (1:5000, ab7291, Abcam), mouse monoclonal antibody against Esrrb, (1:1000, PP-H6705-00, Bio-Techne) rat monoclonal antibody against Sox2, (1:500, 14-9811-80, eBioscience), and rabbit monoclonal against vinculin (1:3000, ab19002, Abcam). The membrane was washed and the indicated horseradish-peroxidase conjugated secondary antibodies were applied: horse anti-mouse IgG (1:5000, Cell Signaling) and goat anti-rabbit (1:5000, Cell Signaling) and goat anti-rat HRP conjugated (1:5000, SC2032, Santa Cruz). Detection was achieved using ECL prime Western blotting Detection system (RPN2133, GE Healthcare).

### Immunofluorescence

Cells were cultured in Ibid tissue culture dishes (IB-81156, Ibid) coated with 0.1% gelatin, washed twice with calcium and magnesium free PBS and fixed in 4% PFA for 20min at RT. The cells were permeabilised in 0.5% Triton X-100 (T8787, Sigma) in PBS for 15min at RT. Subsequently, blocking was performed in 5% donkey serum (D9663, Sigma) and 0.1% Triton X-100 for 1hr at RT. The following primary antibodies in blocking buffer were applied for the indicated protein: Sox2, (1:500, 14-9811-80, eBioscience) and β-catenin (1:1000, 4627, Cell Signaling). Primary antibodies were incubated overnight at 4°C. Subsequently excess primary antibody was washed away and anti-rat Alexa Flour 594^®^ conjugated antibody (1:1000, A21209, Abcam) was added for Sox2, and incubated for 1h at RT. Excess secondary antibody was washed away and DAPI (1:1000, D9542, Sigma) was added and incubated for 10min at RT. Cells were washed and mounted in RapiClear (RCCS002, Sunjin lab).

## Acknowledgements

The CAG-Cre puro plasmid was a kind gift from B. Hendrich (WT-MRC Cambridge Stem Cell Institute, UCAM). We thank Masaki Kinoshita for advice regarding antibodies. A.A-R. is supported by MRC, A.M.is supported by Wntsapp, Marie Curie ITN, J.F. is supported by the Wellcome Trust, W.C.S received core grant support from the Wellcome Trust to the Wellcome Trust Sanger Institute. B-K.K. and R.M. is supported by a Sir Henry Dale Fellowship from the Wellcome Trust and the Royal Society [101241/Z/13/Z] and receives core support grant from the Wellcome Trust and MRC to the WT - MRC Cambridge Stem Cell Institute.

## Author contributions

A.A-R., WC.S. and B-K.K. wrote the manuscript. A.A-R, J.F, WC.S. and B-K.K. designed the FLIP cassette targeting vector. A.A-R., A.M., R.M., and K.A. performed the experiments. S.P. and T.G. performed the bioinformatics analysis. WC.S. and B-K.K. supervised the project.

## Competing financial interest

The authors declare no financial interest.

